# The lysogenization of the non-O157 *Escherichia coli* strains by stx-converting bacteriophage phi24B is associated with the O antigen loss and reduced fitness

**DOI:** 10.1101/860106

**Authors:** A.K. Golomidova, A.D. Efimov, E.E. Kulikov, A.S. Kuznetsov, A.V. Letarov

**Affiliations:** Winogradsky Institute of Microbiology RC Biotechnology RAS, Moscow, Russia; Faculty of Biology, Lomonosov Moscow State University, Moscow, Russia; Phystech School of Biological and Medical Physics, Moscow Institute of Physics and Technology, Moscow, Russia

**Keywords:** stx-converting bacteriophages, shiga-toxigenic *E. coli*, LPS, O antigen, bacteriophage adsorption, temperate bacteriophages

## Abstract

The ability of the Shiga-toxigenic *E. coli* (STEC) to produce the toxin depends on the lysogenic conversion by stx-bacteriophages. The canonical stx-phage phi24B can lysogenize a wide variety of *E. coli* strains. *In vivo* the secondary lysogenization of symbiotic *E. coli* strains by the phages released by infecting STEC populations may contribute to the overall patient toxic load and to lead to the emergence of new pathogenic STEC strains. However, in our experiment all the phi24B lysogens obtained from the environmental *E. coli* isolates had compromised O-antigen (Oag) biosynthesis. These lysogenic strains gained the sensitivity to the T5-like bacteriophages and featured increased sensitivity to the bactericidal activity of the horse serum. We conclude that in most of *E. coli* strains the Oag effectively restricts phi24B infection. The lysogenic clones predominantly rise from the Oag deficient mutants and therefore they have reduced fitness compared to the parental strain.

## 1. Introduction

The verotoxigenic (VTEC) and shigatoxigenic (STEC) *Escherichia coli* strains are associated with multiple foodborne diseases causing morbidity and mortality in humans^1-3^. VTEC and STEC are zoonotic pathogens usually transmitted in the agricultural loop^4^. The majority of STEC strains belong to the O157:H7 serotype^5^, although non-O157 STEC strains have been identified and currently gain increased importance^3^ as, for example, the so-called “Big Six” - O26, O45, O1; O111,O121 and O145^6^, as well as the O104:H4 serotype that caused the well-known 2011 outbreak in Germany^7^.

STEC strains possess a number of pathogenicity factors, the foremost being Shiga toxin production^5, 8^.

Although the Stx-converting bacteriophages are quite divergent genetically and morphologically, from the genome organization perspective all of them belong to a broad group of lambdoid bacteriophages^9, 10^. In these phages, the toxin gene *stx* is located downstream of the conserved gene Q encoding the antiterminator of the late gene region^9^. Toxin expression is repressed in normally growing lysogenic bacterial cells, and takes place only upon the prophage induction. Toxin molecules lacking the signal leading peptide for secretion are released upon cell lysis. The lysogeny in stx-converting phages is less stable compared to stx-lambdoid phages^11-14^ resulting in higher rate of spontaneous induction and in increased sensitivity to environmental factors. Many antibiotics also increase the induction rate of Stx-converting prophages thus enhancing the toxin production. Therefore, the use of antibiotics to treat STEC infections remains controversial^15^.

At the same time, the STEC infections are self-limiting, and the pathogen gets spontaneously eliminated in ca. 2 weeks. The standard for the treatment of these infections relies on supportive care (symptomatic treatment, plasma exchange, infusion therapy) aiming at stabilization of the patient condition during the time required for self–curing of the infection^16^.

Thus it is possible to speculate that the severity of the symptoms and the outcome of the disease may also depend on interaction of the stx phage released by the STEC population in the upper intestine with the resident *E. coli* population in the hindgut. In case of active phage multiplication in this site, the released toxin may contribute to the overall toxin load. However, stx phages are seldom able to form plaques *in vitro* on isolated symbiotic gut *E. coli* strains^17^.

About 70% of Stx-converting bacteriophages are podoviruses related to the bacteriophage vb_EcoP_24B, also known as phage ϕ24B^14, 18^. The phage ϕ24B lysogenization host range was reported to be much broader than its range of hosts that support plaque formation^17^. The same observations were also made for some other stx phages^19, 20^. The establishment of the lysogenic *E. coli* population in the patient’s hindgut may also represent a threat of inducible increased toxin load. The route of lateral toxin gene transmission to other (potentially) enteropathogenic *E. coli* strains adapted to gut environment may lead to emergence of new highly virulent STEC lineages^3, 19^.

The secondary (terminal) receptor of bacteriophage ϕ24B has been identified as BamA protein, previously referred to as YaeT^21^, responsible for insertion of the newly synthesized beta-barrel outer membrane proteins into the bacterial outer membrane^22^. BamA protein is essential for bacterial cell viability and is therefore highly conserved. This circumstance allows speculating that a large variety of the non-Stx-producing or even non-pathogenic *E. coli* strains can be potentially lysogenized *in vivo* and thus get involved in STEC evolution and/or pathogenesis of the STEC-induced diseases.

The available data suggest that the presence of a suitable secondary receptor is not the only factor required for successful phage adsorption and DNA delivery into the host cell. For *E. coli*, it has been shown that many O-antigen types protect the cells nearly completely against the phages not able to recognize O-antigen specifically^23-26^. This is achieved by non-specific shielding of the intimate cell surface by this structure. It was unclear how phage ϕ24B and related viruses that encode only one potential tail spike protein, gp61^14^, may penetrate the O antigen shield in diverse *E. coli* strains belonging to different O-serotypes.

It is possible to speculate that, in the experimental conditions used by James et al.^17^, when a massive amount of the phage is added to the host cell suspension, some phage particles may by chance penetrate through the hypothetical temporary brakes that may exist in the O-antigen shield of some cells. This will be enough to detect a certain frequency of lysogenization, especially if a phage marked with an antibiotic resistance gene is used. Alternatively, the phage may lysogenize the small fraction of the mutant cells depleted of the O-antigen biosynthesis that is normally present in bacterial cultures^25^.

The bacteriophages that are potentially able to infect the strain but are restrained by its O-antigen can be successfully used as a probe for testing the efficacy of the O-antigen-mediated protection^25^. We developed the use of a T5-like bacteriophage DT571/2 mutant lacking lateral tail fibers (LTFs) as such a probe^24^. This phage is hereafter referred to as FimX. Alternatively, the physical state of O-antigen of a culture can be directly assessed by rapid LPS profiling based on SDS-PAGE electrophoresis with sugar-specific silver staining^25^.

## 2. Material and methods

### 2.1 E. coli and bacteriophage strains and their cultivation

The *E. coli* strain MG1655 lysogenized for phage ϕ24B:cat was a kind gift of Prof. G. Wegrzyn, University of Gdansk, Poland. Phage T5 was a gift of Dr. V. Ksenzenko (Institute of protein research RAS, Puschino-na-Oke, Russia). We previously described T5-like bacteriophages of DT57C species and their LTF mutants^24^. These include: phage DT57C, phage DT571/2, DT571/2 ltfA^−^ mutant lacking the LTFs (hereafter FimX) and DT571/2 mutant ABF that carries LTF non-branched LTF with only one receptor-binding domain (instead of two such domains on the branched LTFs of the phages DT57C or DT571/2). Bacteriophage 9g, a siphovirus representing the type strain of the genus Nonagvirus ^27^. Gostya9 is a T5-like bacteriophage that was shown to recognize a different secondary receptor distinct from the receptors of the phages T5, DT57C and 9g ^28^. Bacteriophage G7C, a N4-related podovirus specifically recognizing O antigen of E. coli 4s strain was isolated and characterized by us previously ^29, 30^. We isolated all the above-mentioned phages except for T5 and engineered phage mutants from horse feces as it described in the corresponding publications cited above. The wild *E. coli* strains were previously isolated by us from horse feces and characterized. These were 4s (O22)^23^, HS1/2 (O87)^31, 32^, HS3-104 (O81)^33^, F5 (O28 ab)^34^, and F17 (new O-serotype)^35^. The clinical uropathogenic *E. coli* isolates UP1 and UP11 were received from the clinical microbiological facility of the Institute of Epidemiology (Moscow, Russia). UP11 strain was further identified as an O5 O-antigen producer^36^.

The ability of the strains to produce O antigens was controlled by LPS profiling as described in Kulikov et al. (2019)^25^.

*E. coli* 4s and F17 rough variants 4sR (a *wclH* mutant of 4s)^23^ and F17 *wbbL*^−35^ were engineered by us previously.

All the *E. coli* strains were cultured on LB medium (trypton 10 g, yeast extract 5 g, NaCl – 10 g, distilled H_2_O – up to 1 l). This medium was supplemented with 15 g of bacto-agar per 1 l for plates or with 6 g of bacto-agar for top agar.

Bacteriophage FimX was propagated on E. coli 4sR and enumerated using the conventional double-layer plating technique.

Bacteriophage ϕ24B:cat was obtained by mitomycin C induction of *E. coli* MG1655 (ϕ24B:cat) strain. For this procedure, the overnight culture of the lysogen was grown in the presence of 34 μg/ml of chloramphenicol. Then 300 ml of LB in 500 ml Erlenmeyer flask was inoculated with 3 ml of the overnight culture (N.B. – this volume ratio gave a better phage yield than conventional conditions with better aeration). The culture was grown in the orbital shaker at 220 rpm, 37°C up to OD_600_ = 0.2. The mitomycin C was then added up to 1 μg/mL and the incubation was continued overnight at the same conditions. After the incubation, lysis of the culture was observed. The lysate was cleared by centrifugation at 15000 × g for 15 min. The supernatant was collected, PEG-precipitated^37^, and resuspended in 3 mL of SM buffer (Tris-HCl pH 7.5 – 10 mM, NaCl – 50 mM, MgCl_2_ – 10 mM, gelatin – 5 g/l). The phage stock was titered and used in these experiments.

For titration of the phage ϕ24B:cat, a modified double-layer technique was used. The top-layer medium contained 4 g/l of the bacto-agar (instead of 6 g/l) and was supplemented with CaCl_2_ up to 5mM. The bottom layer was supplemented with 2.5 μg/ ml of chloramphenicol. 300 μg of log-phase culture of *E. coli* C600 (OD_600_ = 0.6) was used for the lawn inoculation.

### 2.2 Lysogenization of the E. coli strains

This procedure was performed as described in James et al.^17^ with minor modifications. Briefly, a mid-log liquid culture of an appropriate strain was grown in LB medium, the phage was added at a multiplicity of 5 pfu/host cfu, and the mixture was incubated at 37°C for 30 min. After the incubation, the cells were spun down in a table-top centrifuge (10000 × g, 1 min), the cells were resuspended in LB, washed twice with LB to remove non bound phage and plated on plates supplemented with 34 μg/ml of chloramphenicol for lysogen selection.

### 2.3. LPS profiling

by SDS-PAGE electrophoresis was performed as recently described ^25^.

### 2.4 Serum bactericidal activity (SBA)

against different strains was measured as follows. The blood samples collected from clinically healthy horses for routine veterinary control purposes were used. The samples were collected into a yellow-cap vacuum tube with clot activator (Elamed, Moscow, Russia). The serum was separated by centrifugation at 1600 × g for 10 min. The serum was stored at +4 °C and used for the tests within 24 h. For the SBA assessment, the wells of 96-well plate containing 175 μl of LB medium and 25 μl of the serum were inoculated with 5 μl of the corresponding strain mid-log phase culture (OD_600_ = 0.6) and the plates were incubated at 37°C in an automated plate reader with agitation. The OD_600_ was recorded every 30 min. In the control experiment the same volume of physiological saline replaced the serum. The whole experiment was triplicated.

## 3. Results

We decided to use these two approaches to evaluate the O antigen production status of the ϕ24B lysogens generated in environmental *E. coli* isolates. To do so, liquid cultures of the O antigen producing strains 4s, HS1/2, HS3-104, F5, F17, UP1 and UP11 and of the rough strains 4sR and C600 were challenged with phage ϕ24B:cat as described by James^17^. The lysogens were then selected by plating the mixture on LB plates supplemented with 34 μg/ml of chloramphenicol. The lysogens were obtained for strains 4s, HS1/2, HS3-104, F5 and F17. No lysogens were observed on strain UP11. The lysogenization frequency was about 10^−4^ lysogen cfu/ phage pfu for the rough strains and about 10^−7^ – 10^−6^ in O antigen-producing strains. The latter value is comparable to the level of spontaneous mutations in *E. coli* inactivating a medium-sized gene (e.g. phage-resistant mutants).

We selected 3 lysogen clones per strain and confirmed the ϕ24B prophage presence using PCR for gene 61 (the tailspike protein gene). For *E. coli* 4s lysogens we also performed mitomycin C induction followed by transmission electron microscopy that confirmed that a phage morphologically identical to ϕ24B was produced.

LPS profiling of the lysogens obtained indicated that in all cases these strains did not produce O-antigen at all or the O-chain synthesis was greatly decreased compared to the parental strains (Fig. 1).

**Figure 1.**
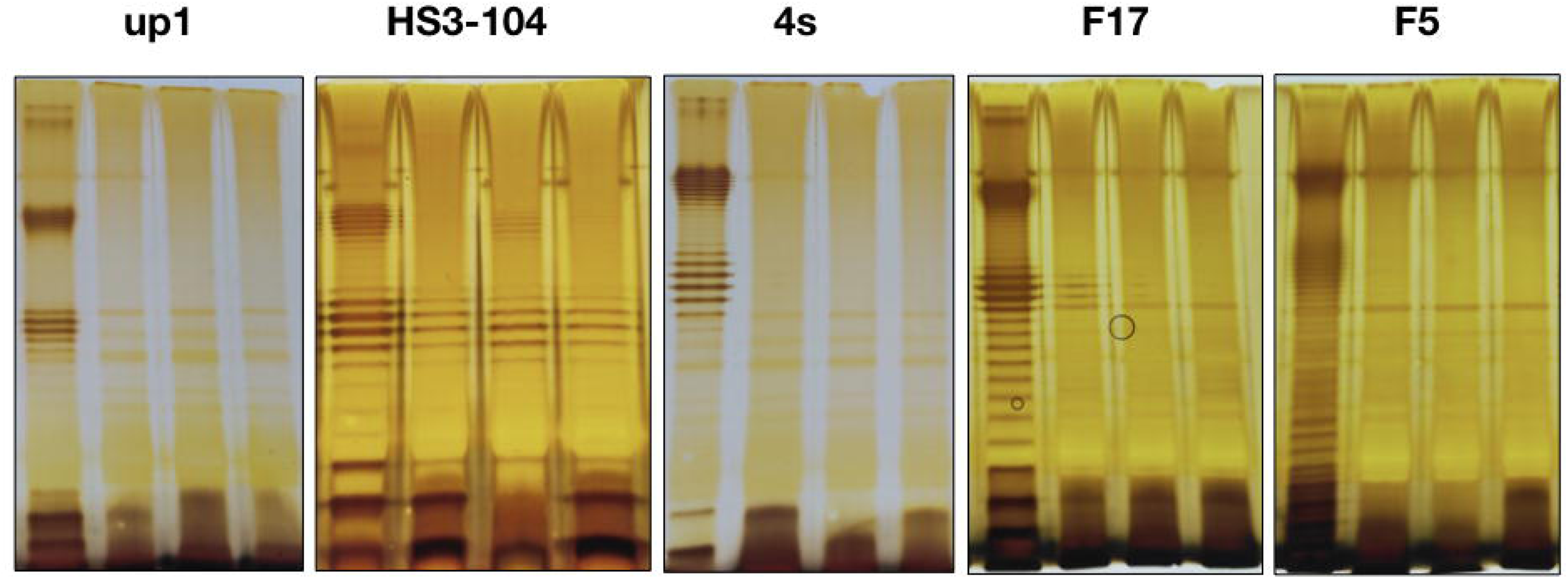
LPS profiles of the *E. coli* strains used and their derivative ϕ24B:cat lysogens. The left lane on each of the panels – the wild type cells, other lanes – three lysogenic clones for each strain.

We tested the ability of phage FimX to grow on the lawns of the lysogens obtained. This phage was not able to form plaques on the parental O-antigen – producing strains, except for F5 on which it formed plaques with an efficiency of plating (EOP) of 10^−4^ compared to the C600 strain used for FimX propagation ^34^. At the same time the EOP of FimX phage on all the lysogenic cultures tested was in the range of 0.1 – 1.0 compared to the *E. coli* C600 strain. The effect of lysogenization on FimX growth was not distinguishable from other methods of rough mutant generation previously used by us in 4s or F17 strains^23, 25^.

The other T5-like phages (DT57C, DT571/2, ABF and Gostya9) as well as the siphovirus 9g demonstrated the gain of the infectivity on the lysogens derivatives of some strains that were initially resistant to these phages. Phage G7C that is dependent on the specific O antigen recognition for infection of E. coli 4s cells^30^ was not able to infect *E. coli* 4s (ϕ24B:cat) lysogenic strains in good agreement with O antigen production loss detected by the LPS profiling (Fig. 1).

Since the O antigen synthesis compromised strains are believed to be more vulnerable to immunity factors, we decided to measure the susceptibility of the lysogens obtained to the bactericidal activity of the horse serum (SBA). All the wild type strains were resistant to SBA in our conditions. Their cultures grew in presence of the serum as well or even slightly more rapidly than in the control experiment. In the absence of the serum the lysogenic strains showed the growth rates close to their cognate wild type strains. At the same time the growth of the lysogens was almost completely abolished in the presence of the serum (Fig. 2). Only one of the lysogenic clones tested, the derivative of the strain HS3-104, was able to grow significantly in presence of the horse serum, though the rise of the optical density was delayed and the growth rate was significantly lower than in the parental strain (Fig.2). This result can be explained by the fact that in HS3-104 lysogens the O antigen synthesis was strongly decreased but not completely abolished (Fig.1). So, the actual synthesis of O-polysaccharide could be upregulated in this particular clone in the conditions of the experiment.

**Figure 2.**
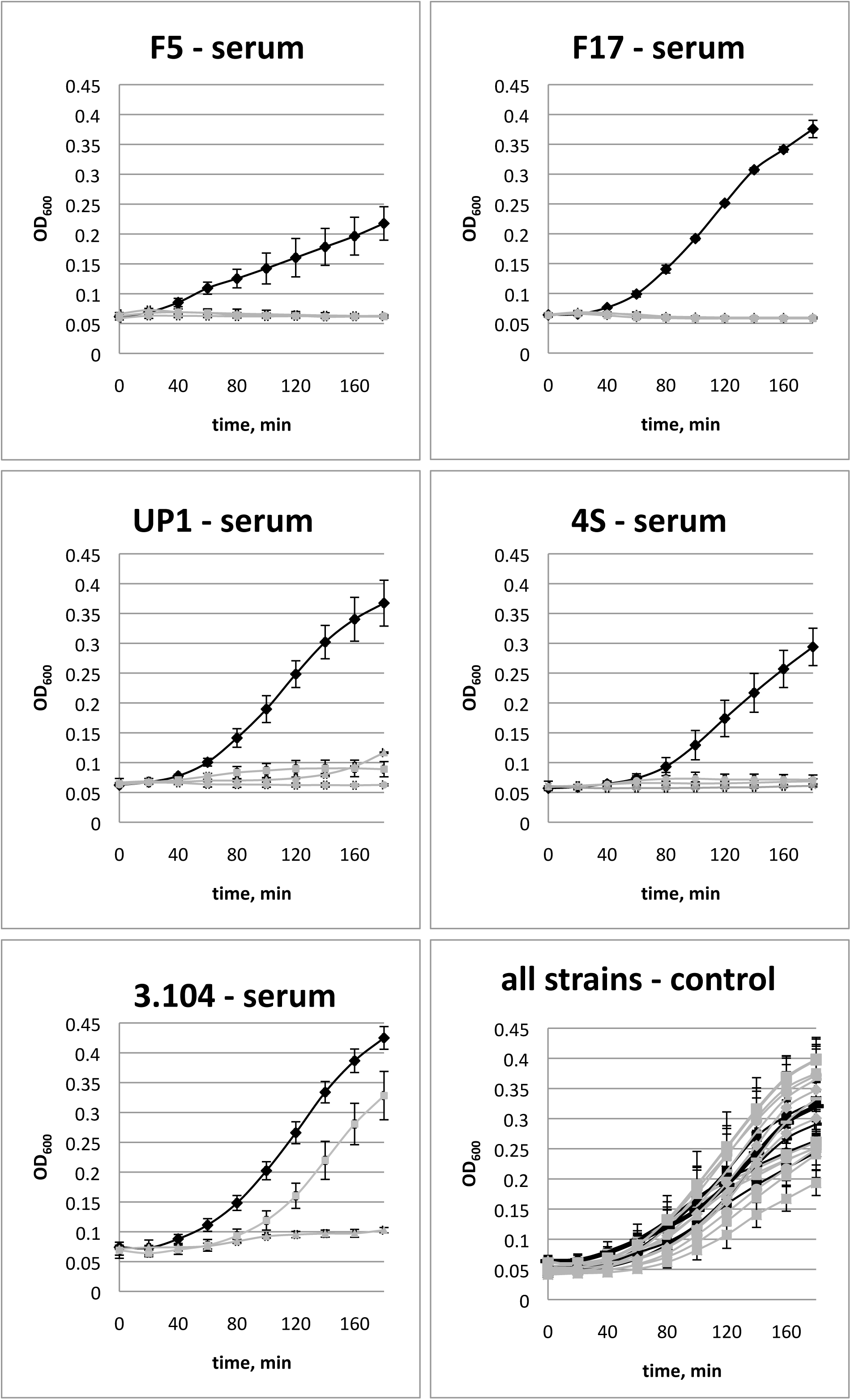
Sensitivity of the *E. coli* strains and their derivative ϕ24B:cat lysogens’ growth to the horse serum bactericidal activity. Black lines – the wild type strain, grey lines – three lysogenic clones tested for each original strain.

## 4. Discussion

The results obtained allow us to conclude that lysogenization by the phage ϕ24B of diverse *E. coli* strains producing O antigens was not due to an unusual ability of this virus to penetrate the O antigen shield, but was mediated by spontaneous formation of bacterial rough mutants or of mutants with significantly compromised O antigen biosynthesis. It is not clear why the lysogenization was not effective for some strains. The activity of antiviral systems, such as restriction-modification, avoiding the lysogenization at stages after the viral DNA penetration into the cell^38^, cannot be excluded. Also the effect may be due to point mutations present in BamA protein or lower frequency of rough mutants in particular strains.

In the conditions of our experiment, the high concentration of bacteriophage used allowed almost all the cells potentially susceptible to the phage to be infected. However, *in vivo* the populations of *E. coli* are very unlikely to face such a massive viral attack. The fraction of rough mutants in natural habitats is hard to estimate, but we can speculate that it should be lower than in *in vitro* conditions because such mutants have compromised protection not only from the phage attack but also from immune system agents such as serum bactericidal activity^39-41^ and from other environmental factors^26^ and therefore should be counter-selected. Moreover, if stx-phage lysogens were formed by infection of such rough mutants, their expected fitness and/or virulence would be significantly lower than that of the parental strains. These strains, noteworthily, were highly sensitive to SBA of the horse serum to which the parental O-antigen producing strains were completely resistant. Therefore the lysogens for the ϕ24B phage are expected to have reduced virulence. Thus, the factor of non-specific protection of the bacterial cells by the O antigen should not be neglected during the evaluation of the potential significance of stx-converting phage transmission in nature (as it currently is neglected in many studies^19, 20^).

We also should note that the lysogenization by ϕ24B:cat appears to be a simple and efficient procedure for selection for mutants with compromised or completely abolished O antigen synthesis. This procedure may be particularly valuable for the researchers working with field isolates of *E. coli* for which the genomic sequences are not yet available and/or in which other rapid techniques such as recombination with PCR fragments for genes knockout^42^ are frequently less effective than in laboratory *E. coli.*

## Acknowledgements and funding

We are grateful to Dr. S. Bloch, Dr. B. Nejman-Faleńczyk and Prof. G. Wegrzyn from the University of Gdansk, Poland for their help in establishing the procedures of phage ϕ24B cultivation in our lab and to Dr. E. M, Kutter from the Evergreen state college, Olympia, USA, for critical reading and linguistic correction of the manuscript.

This work was supported by the RSF under Grant 15-15-00134P.

---

**Table 1.** Sensitivity of the E. coli strains and their derivative ϕ24B:cat lysogens to virulent coliphages.

|  | **E. coli strains** |  |  |  |  |  |  |  |  |  |
| --- | --- | --- | --- | --- | --- | --- | --- | --- | --- | --- |
|  | **HS3-104** |  | **4s** |  | **F5** |  | **F17** |  | **UP1** |  |
| **Phages** | **wt** | **Lyso-gens** | **wt** | **Lyso-gens** | **wt** | **Lyso-gens** | **wt** | **Lyso-gens** | **wt** | **Lyso-gens** |
| **DT57C** | + | + | + | + | +/- | + | - | + | - | + |
| **DT571/2** | + | + | - | + | +/- | + | - | + | - | + |
| **fimX** | - | + | - | + | +/- | + | - | + | - | + |
| **ABF** | + | + | - | + | +/- | + | - | + | - | + |
| **Gostya9** | - | + | - | + | + | + | - | + | - | - |
| **G7C** | - | - | + | - | - | - | - | - | - | - |
| **T5** | - | + | - | + | - | - | - | + | - | + |
| **9g** | - | + | - | + | + | + | + | + | - | + |

+/- – very small turbid plaques with EOP < 10<sup>-3</sup> in respect to plating on the optimal host strain

